# Joint Alignment and Tree Inference

**DOI:** 10.1101/2021.09.28.462230

**Authors:** Jūlija Pečerska, Manuel Gil, Maria Anisimova

## Abstract

Multiple sequence alignment and phylogenetic tree inference are connected problems that are often solved as independent steps in the inference process. Several attempts at doing simultaneous inference have been made, however currently the available methods are greatly limited by their computational complexity and can only handle small datasets. In this manuscript we introduce a combinatorial optimisation approach that will allow us to resolve the circularity of the problem and efficiently infer both alignments and trees under maximum likelihood.

## 1. Introduction

In this manuscript we introduce a hill-climbing combinatorial optimisation approach for tree and multiple sequence alignment (MSA) inference under the Poisson Indel Process (PIP) [1] model of sequence evolution. Our goal is to produce maximum likelihood (ML) estimates simultaneously for the tree and the MSA. We aim to resolve the circularity of MSA and tree inference, as most of the existing methods need a guide tree to reconstruct an MSA from unaligned sequences, while tree reconstruction methods require an MSA as input.

Moreover, it has been previously shown that standard ML analyses that either ignore gaps or treat gaps as missing data may lead to statistical inconsistency when gaps are present in the alignment [2]. We avoid that inconsistency by using a sequence evolution model that appropriately accounts for gaps. PIP models sequence evolution not only by substitutions, but also by insertions and deletions (indels) and defines the proper joint likelihood of the MSA and the phylogenetic tree with indels.

Some existing works reconstruct trees and MSAs simultaneously using evolutionary indel models in the Bayesian framework [1, 3]. Unfortunately, these methods are greatly limited by their computational complexity and consequently by the amount of computation necessary to produce reliable estimates. For example, [1] tested their method on 7 taxa trees, whereas in [3] the baseline dataset size is 5 taxa, while the method is capable of handling datasets of up to 12 taxa. Unfortunately, such dataset sizes are rarely enough for scientifically informative analyses. In particular, epidemiological studies may need to effectively handle from several hundred up to thousands of sequences.

We opt for the frequentist approach in our implementation which would reduce the computational complexity and would allow us to process bigger datasets. We base our solution on an established approach to ML tree search, implemented in PhyML [4], and on efficient dynamic programming alignment under PIP [5]. The basic idea is to improve the starting tree given an initial alignment, similar as to what PhyML does for fixed alignments in general, and then improve the global likelihood of the tree by optimizing the alignment given the updated tree structure, repeating the two improvement steps until convergence.

## 2. Methods

### 2.1. Overview

We propose a way to estimate both the phylogenetic tree and the MSA using two-phase combinatorial optimisation. In the first phase we propose to do tree search using subtree pruning and regrafting (SPR) moves, applying several moves at a time to get to a new tree with an improved likelihood based on a fixed MSA. In the second phase, we propose to iteratively realign the sequences on the modified tree to improve the fit of the MSA given the new tree structure. We propose a way to realign the sequences guided by global likelihood values, therefore guaranteeing that the new MSA typically increases, but never decreases, the likelihood on the modified tree.

### 2.2. Tree search

We base our tree search on the most basic version of PhyML 3.0 SPR approach as described in [4]. In PhyML, the tree search starts with an initial tree, which can either be a tree built with a fast distance-based method, e.g. BioNJ, or a random tree on the given taxa. Given the initial tree, the free parameters of the sequence evolution model are adjusted to increase the likelihood of the starting phylogeny using a numerical optimisation method. After the initial adjustments, PhyML iteratively adjusts the phylogeny and model parameter values such that the likelihood increases with each iteration. The tree is adjusted by simultaneously applying a set of SPR moves that each individually improve the tree likelihood given the current parameter values. Then the parameters of the evolutionary model are adjusted, and the branch lengths of the new phylogeny are optimised. Finally, the likelihood for the new tree and model parameter combination is computed and tested against the value from the previous iteration, and the cycle ends when no significant improvement is found.

Conceptually, as in PhyML, per iteration of combinatorial optimisation we will evaluate the tree likelihood for each possible SPR move and will select a subset of the moves that increase the tree likelihood. The function `Multiple_SPR_Cycles`(*τ,* Θ) (Algorithm 1) is defined similarly as to how [4] define tree search. We propose to also include an MSA update step after each movement in the tree space (Algorithm 1, line 9).

#### Algorithm 1

The tree search algorithm pseudocode given by the `Multiple_SPR_Cycles` function, an adapted version of the function with the same name from [4].

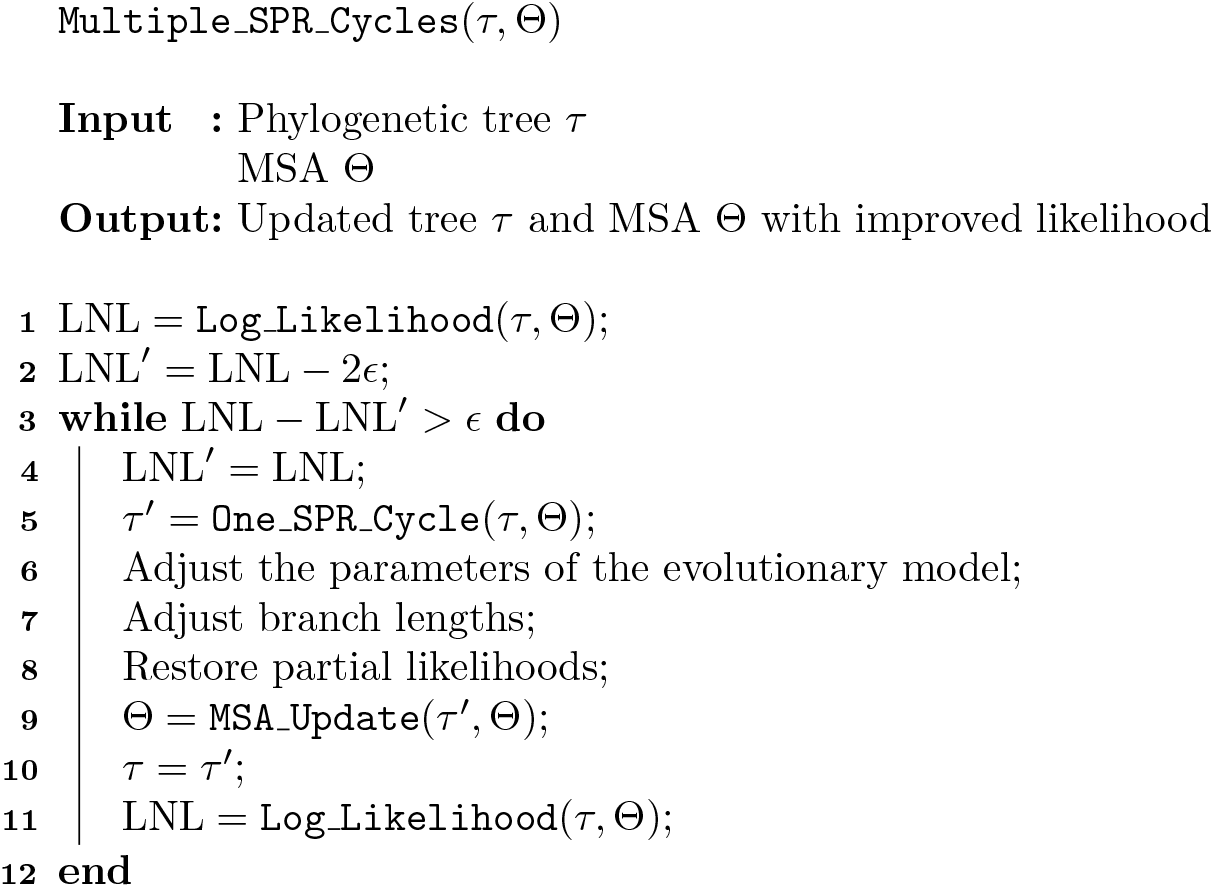

#### 2.2.1. Tree likelihood under PIP Log Likelihood(*τ,* Θ)

The PIP model of molecular evolution uses a Poisson process and an augmented continuous-time Markov chain formulation of the substitution process to describe single nucleotide insertion, deletion and substitution events on the tree [1]. PIP models insertions as a Poisson process on the tree meaning that the insertion rate does not depend on sequence length. This gives PIP a significant computational advantage over classical indel models such as TKF91 [6] and TKF92 [7] (marginal likelihood in exponential time), as the marginal likelihood of a tree under PIP can be computed in linear time.

PIP defines a string-valued evolutionary process along the branches of a binary rooted phylogenetic tree *τ* = (*V, E, b*) on *N* leaves, where *V* is the set of 2*N* − 1 tree nodes, including the set of leaves *L* ⊂ *V*, and the set of edges *E* ⊂ *V* × *V*. The set of leaves *L* includes all observed sequences represented as strings of characters from a finite alphabet Σ (e.g. for nucleotides, Σ = {`T, C, A, G`}) The root node of the tree Ω ∈ *V* is the most recent common ancestor of all leaves. For each node, we label *b*(*υ*) the length of the branch leading to node *υ* from its parent. The total tree length ||*τ* || is the sum of all branch lengths *b*(*υ*). We label the MSA Θ, where *c* ∈ Θ is a single column of the MSA and |Θ| = *M* is the total number of sites in the MSA.

PIP models atomic insertion events as a Poisson process on the tree with the rate measure

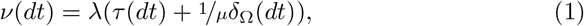

where *λ* is the insertion rate, *μ* the deletion rate, *τ* (·) is the distance from the root to a given point on the tree, and *δ*_Ω_(·) is the Dirac delta function. This insertion rate measure guarantees that the expected sequence length remains constant through time. The deletions and substitutions are modelled by a continuous-time Markov chain on an extended alphabet Σ_*ϵ*_ = Σ ∪ *ϵ*, where *ϵ* denotes a gap. The gap character is an absorbing state, meaning that once a character is deleted it cannot reappear, which ensures that the homology paths start with a unique insertion event. We can select an arbitrary character substitution model with an instantaneous rate matrix *Q* (e.g. HKY85 for DNA or WAG for protein sequences), extended to include the gap character:

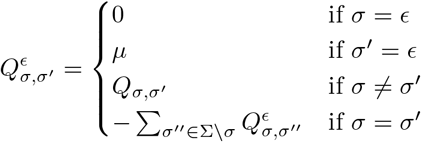

For a node *υ* the probability that a single character is inserted on the branch leading to it is:

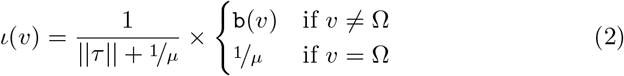

The insertion probability per node is proportional to the length of the branch leading to it, while the root node Ω has a non-zero insertion probability set such 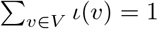.

For a node *υ* the probability that a character inserted on the branch leading to node *υ* will survive until node *υ* is:

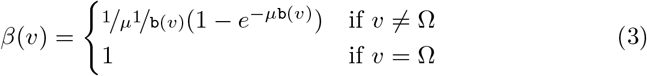

The marginal likelihood of the tree and the MSA under PIP is computed as follows:

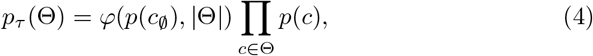

where *p*(*c*) is the marginal likelihood of a single column in the MSA, and *p*(*c*_∅_) is marginal likelihood of an unobserved (empty) MSA column.

The marginal likelihood has to account for an unknown number of empty columns – sites for which the character goes extinct before reaching any of the tree leaves. The number of such columns is potentially unbounded and is unknown. However, these columns are invisible in the final alignment and are interchangeable, which allows us to marginalise over all possible such columns analytically. This marginalisation factors in the probability of randomly sampling an unknown number of evolutionary histories of which we only observe |Θ|, while the rest are empty columns.

The marginalisation is simplified by introducing the function *φ* for all for all *p* ∈ (0, 1) and *m* ∈ {1, 2, …}:

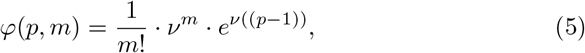

where

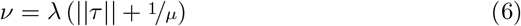

The individual column likelihoods are:

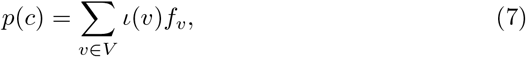

where *f*_*υ*_ is the probability of the homology path explaining column *c* for the subtree rooted at *υ*, if we assume that the character was inserted at *υ*. The factors *f*_*υ*_ can be computed in O(N) using a variant of Felsenstein’s peeling recursion.

Let *S* be the subset of leaves that have a non-gap nucleotide in the column *c*. This subset of leaves defines a set of possible insertion locations – a subset *A* of the internal nodes on the tree that are ancestral to all the leaves in *S*. We can compute *f*_*υ*_ as:

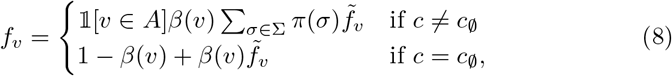

where

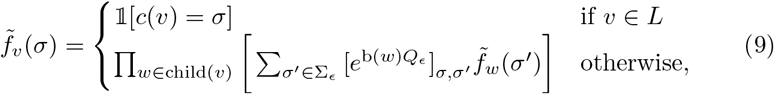

and

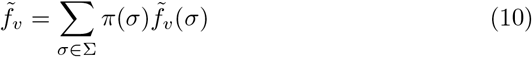

The Log Likelihood(*τ,* Θ) function computes the logarithm of the likelihood *p*_*τ*_ (Θ) of the tree *τ* and PIP given the MSA Θ. The partial likelihoods for the substitution-deletion process 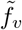 per site can be computed in linear time with respect to the number of leaves on the tree. Similarly, the factors *f*_*υ*_, which account for the insertion probabilities at different nodes, and the sets of ancestral nodes *A* can also be computed in linear time, i.e. the overall computation of the likelihood under PIP requires *O*(*NM*) operations.

#### 2.2.2. *SPR tree move selection and application* One_SPR_Cycle(*τ,* Θ)

The function `One_SPR_Cycle`(*τ,* Θ) here is defined the same way as in [4] Appendix 1, returning the updated tree structure. SPR moves are ranked by the increase in the likelihood of the new tree structure and then several best moves are applied simultaneously (see Figure 1a). However, we want to minimise the number of realigning operations necessary, thus we will only realign the parts of the tree that were affected by any one of the applied moves.

**Figure 1:**
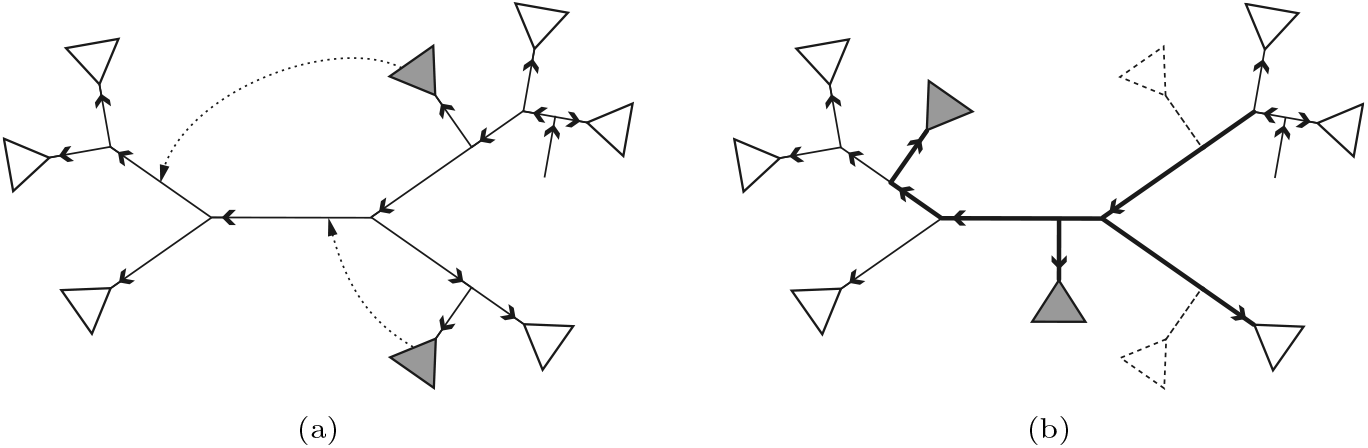
Tree structures representing the current iteration of `One_SPR_Cycle`(*τ,* Θ). The tree is rooted at an arbitrary branch, and branches show the directionality of time flow, which defines partial likelihood values. Figure 1a shows the two best SPR moves selected in `One_SPR_Cycle`(*τ,* Θ), applied simultaneously to update the tree structure. Figure 1b shows the spanning “dirty” subtree of branches that were affected by the moves. It contains pruned branches, branches that lost a node as the result of the pruning, as well as all branches necessary to create a connected spanning subtree.

To do so, we will modify the original `One_SPR_Cycle`(*τ,* Θ) function to label certain branches as “dirty”. Firstly, we label the branches that were pruned and those that lost a node as a result of the pruning. Secondly, we also mark all the branches between nodes belonging to any of the “dirty” branches, such that a connected spanning tree is marked for realignment (see Figure 1b). We will perform realignment operations on the set of “dirty” branches. Thus, we realign on a limited number of branches, effectively excluding the subtrees that were not affected by the change in tree structure due to the moves.

### 2.3. MSA update

After a single SPR move cycle, we proceed to update the MSA to improve the likelihood. This step requires that we have a rooted tree with fixed partial likelihoods 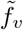 and *f*_*υ*_ (done in line 8 of Algorithm 1) and a connected subtree of all “dirty” branches.

As we are developing a hill-climbing optimisation approach, we need to ensure that any MSA update improves the likelihood of the tree/MSA combination. Note that progressive aligners such as, for example, ProPIP [5], do not align under the global likelihood (likelihood including all sequences) at every step of the process. Instead, during progressive alignment only the taxa within the current subtree are taken into account, and the alignment is evaluated under the likelihood for this particular subtree (local likelihood). All the sequences together are taken into account in the final step of the alignment, when aligning at the root. Progressive alignment is a greedy approach, so the intermediate sub-alignments may not be the best under the global likelihood, however the final alignment of the two sub-alignments at the root necessarily is. We propose to take advantage of global likelihood maximisation at the root and use it to always propose an MSA that improves the likelihood.

We propose to traverse the “dirty” subtree, realigning at every branch. First, we re-root the tree at a dirty branch, adjusting all the partial likelihoods (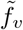 and *f*_*υ*_) to account for the location of the new root. While in general this operation would mean recomputing all partial likelihoods for the new root location, the partial likelihoods correspond to directed branches and remain the same for branches that did not change direction. This means that to initialise the traverse we will have to recompute the partial likelihoods for branches on the shortest path between the old root node and the new root, while the rest remain unchanged, for which the computational complexity is bounded by O(NM). Moreover, this can be done in `Multiple_SPR_Cycles` when restoring the partial likelihoods to account for the updated branch lengths and parameters of the evolutionary model. Once the new root and the partial likelihoods are set up, we propose to realign the homology paths defined by the left and right subtrees under the root, given the current MSA. Essentially, this is the final alignment step defined by the progressive alignment approach under PIP [5].

To realign fully we propose to then visit all the “dirty” branches in order. We move the current realignment root over to a neighbouring dirty branch, updating the partial likelihoods corresponding to the affected branches (see Figure 2a). As with this operation we only change the directionality of one branch, the update can be done in linear time w.r.t. the number of sites and in constant time on the number of tree nodes (O(M)). We call this function `Move_Root_By_One` in the following pseudocode.

**Figure 2:**
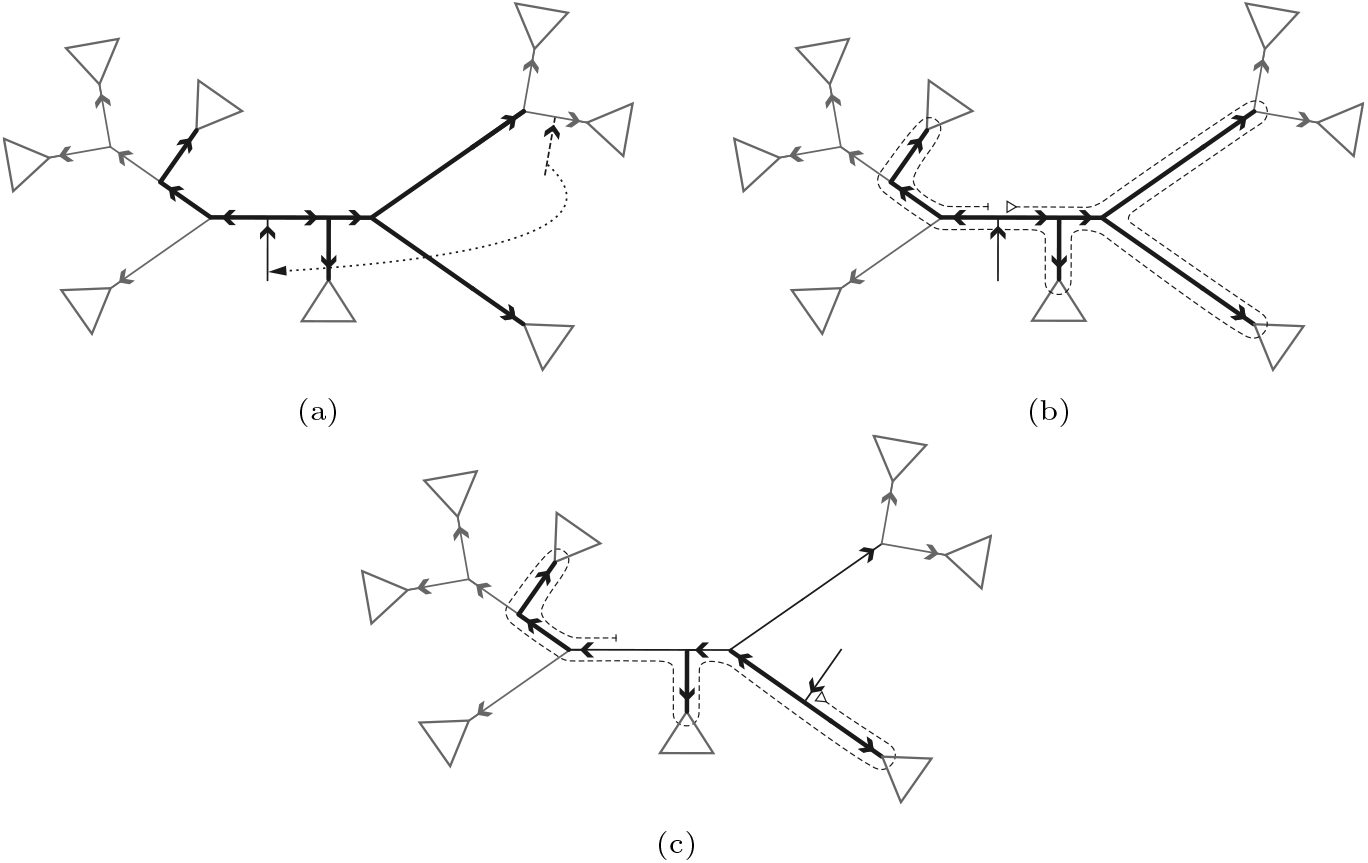
Figure 2a shows relocating the root to within the “dirty” subtree marked by thicker branches to initiate realignment. The branch directionalities change to reflect the change in time flow and the partial likelihoods that need to be recomputed. Figure 2b shows the starting Euler tour through the tree, defining the branch order for realignment. The branch order is based on the initial root location and stays unchanged even though the root is moved to realign at the next branch. Figure 2c shows an intermediate step in the realignment on the whole tree, following the order defined by the Euler tour. The branches that have already been realigned at and lost the “dirtiness” (e.g. the initial root branch) will not be realigned at again, but will be traversed to access the remaining “dirty” branches in the tree.

In summary, we propose to initialise the MSA update by moving the root to a “dirty” branch in our tree. Then we traverse the “dirty” subtree, realigning the two subtrees defined by the current root position and moving the root to a neighbouring “dirty” branch that awaits realignment (Figures 2b and 2c). Finally, we should have traversed all “dirty” branches in the tree and realigned with the root placed at each one of these branches, resulting in an MSA that improves the total tree likelihood. The pseudocode of the proposed approach is shown in Algorithm 2.

#### Algorithm 2

The MSA update algorithm pseudocode given by the MSA Update function.

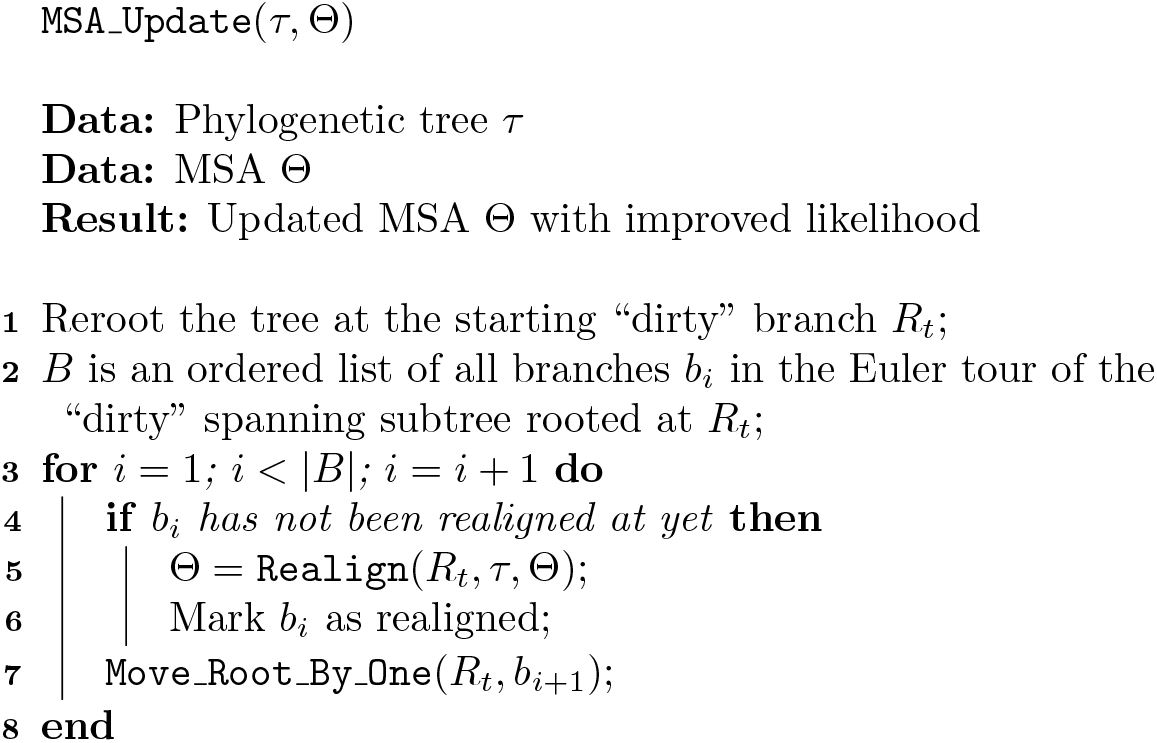

#### 2.3.1. *Moving the root node by one branch* Move_Root_By_One(*R*_*t*_, *b*)

`Move_Root_By_One` is a function that moves the tree root to a neighbouring branch. The realignment root is always placed in the middle of a branch and can move onto any of the four adjacent branches as defined by the unrooted representation of the tree.The prerequisite for the move is that all the intermediate values required to compute the tree likelihood (the partial likelihoods of the substitution-deletion process 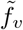, partial likelihoods including the insertion probabilities *f*_*υ*_ and the ancestral sets *A*, defined in section 2.2.1) are valid for the current root position. When moving the root to an adjacent branch, the only values that need to be updated are the values associated with the two affected branches. This update can be done in linear time w.r.t. the length of the MSA and is independent of the number of nodes in the tree, as the rest of the intermediate values remain valid. This move operation is roughly equivalent to rotating the tree, where the pivot point is the center of the branch which we are moving the root to.

#### 2.3.2. *Realigning* Realign(*R*_*t*_, *τ,* Θ)

Realign is a function that realigns the two sub-MSAs defined by the two subtrees of the current root node. We base this MSA adjustment step on the dynamic programming (DP) alignment approach described in [5]. The DP alignment takes the homology structures of the two child subtrees of a node (*R*_*t*_ in this case) represented by the current MSA and by the intermediate values 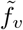 and *f*_*υ*_, and realigns them by maximum likelihood in polynomial time. This way, we realign under the global likelihood values, as the likelihood computed at the root will take into account the full tree structure by realigning at the current root node. Additionally, we reuse the homology paths defined by the already existing MSA, thus not discarding results from the previous steps of the operation, but rather improving the already existing solution.

To get the maximum likelihood alignment of the two sub-MSAs using the DP aligner under PIP, we need to define three score matrices accounting for different intermediate states (column match, gap in one alignment and gap in the other alignment).Traditional sequence alignment under scoring functions that increase monotonically in the alignment length can be solved with two dimensional scoring matrices. As detailed out in [5] the marginal likelihood under PIP is not monotonic. As as consequence, we need to represent the alignment length along a third dimension.

Assuming that the alignments are labelled *X* and *Y*, and each contain *N*_*X*_ and *N*_*Y*_ sequences and |*X*| and |*Y*| characters, we introduce three three-dimensional matrices *S*^*M*^, *S*^*Y*^ and *S*^*X*^ of size (|*X*|+1)×(|*Y*|+1)×(|*X*|+|*Y*|+1) described as follows:

- Match cells: 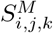 contains the likelihood of the partial optimal MSA of length *k* of the first *i* elements of *X* and the first *j* elements of *Y*, where the columns *Y*_*i*_ and *X*_*j*_ are aligned.
- GapX cells: 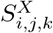 contains the likelihood of the partial optimal MSA of length *k* of the first *i* elements of *X* and the first *j* elements of *Y*, where the column *X*_*i*_ is aligned with a gap in sequence *Y*.
- GapY cell: 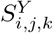 contains the likelihood of the partial optimal MSA of length *k* of the first *i* elements of *X* and the first *j* elements of *Y*, where the column *Y*_*j*_ is aligned with a gap in sequence *X*.

The algorithm works in two phases, forward and backward. In the forward phase, the matrices *S*^*M*^, *S*^*Y*^ and *S*^*X*^ are filled out with values of the corresponding likelihoods, defined as follows:

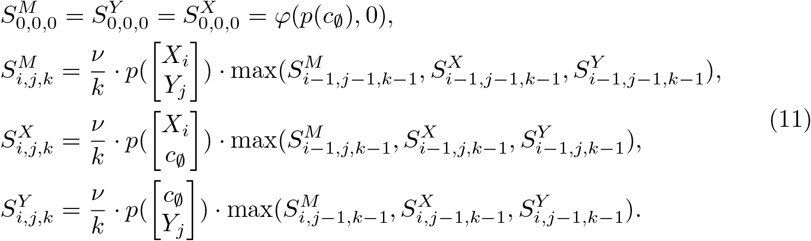

for *i* = 1, …, |*X*|, *j* = 1, …, |*Y*| and *k* = 1, …, |*X*| + |*Y*|. The respective column likelihoods needed are computed in constant time w.r.t. the number of nodes and characters, since all the intermediate values (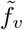, *f*_*υ*_ and *A*) are already computed for the current root position. Consequently, the asymptotic runtime of this phase is *O*(*M*^3^). During the forward phase we maintain a back tracking matrix to record for each position i, j, k the name of the DP matrix (either “ SM”, “ SX”, or “ SY”) with highest likelihood at the same position.

In the backward phase, we traverse the backtracking matrix to progressively construct the maximum likelihood alignment for the two sub-MSAs. For a detailed account of the DP alignment algorithm please see [5].

## 3. Discussion

While the approach described here is based on the PhyML tree search heuristic, it is an independent addition to the algorithm rather than something that is tied inherently to the tree search strategy. We selected the PhyML approach as it is an established method with conclusive results. However, as our method is an additional step in inference rather than a modification, we can switch to any other tree search heuristic that may prove better than the current state-of-the-art. As long as the algorithm can provide a set of affected branches or the affected spanning tree, justified by the underlying approach, we can realign sequences efficiently to iteratively improve the likelihood of the tree/MSA combination.

Moreover, our approach allows us to adapt the particular locations where realignment is either necessary or the most efficient at improving the likelihood. Potentially we could realign at each tree node, going up in asymptotic complexity to *O*(*NM*^3^). However, we think that more sparing realignment strategies will work just as well to improve the tree likelihood. Currently we intend to only realign along the spanning tree affected by the tree moves (“dirty” spanning tree), but it may be just as helpful to realign at a smaller subset of the “dirty” branches, or even to only realign at the regraft locations. These strategies will have to be tested empirically on the initial implementation, which is being developed at the time of this manuscript.

The DP alignment algorithm is also in active development and our group has presented work that significantly speeds up the aligner through the use of fast Fourier transform [8] and through parallelisation [9].

Moreover, PhyML uses a parsimony-based filter to quickly judge whether a potential move will lead to a loss in likelihood, and can therefore be discarded without having to compute the true tree likelihood after such a move. We plan to use a similar approach, using a gap-aware parsimony filter to ensure that we do not waste computation on moves that will surely not improve the tree.

